# State-dependent cannabidiol interactions with fentanyl-bound mouse μ-opioid receptor conformations: a three-state molecular dynamics study

**DOI:** 10.64898/2026.08.24.746804

**Authors:** Jim Wager-Miller, Gergo Szanda, Alex Straiker, Kimberley Bosire, Ken Mackie

## Abstract

We published recently that one of the main constituents of cannabis products, cannabidiol (CBD), is an efficacious negative allosteric modulator (NAM) of the µ opioid receptor (MOR1) (Bosquez-Berger et al., 2023). Here, we investigated how the presence of cannabidiol (CBD) is associated with fentanyl (FEN) binding across MOR1 conformations. We performed molecular dynamics simulations of systems containing FEN alone or FEN+CBD in three mouse MOR1 conformational backgrounds: active-like 5C1M, inactive-like 4DKL, and a modeled Morph50 intermediate between the 5C1M and 4DKL conformations. Three independently seeded 200 ns trajectories were analyzed per model and condition (18 trajectories total), with the trajectory treated as the independent unit. Across the matched 0-200 ns window, consensus CBD contacts and CBD-associated changes in FEN contacts were strongly state dependent. Corrected intracellular TM3–TM6 analyses separated the expected active-like, intermediate, and inactive-like backgrounds but did not identify a CBD-associated shift that was consistent across both geometric definitions and all three replicates. Equal-weight replicate-composite density maps preserved both the shared ligand distributions and this between-trajectory variability. These descriptive results support receptor-state-dependent CBD-FEN-MOR1 interactions while emphasizing the limited inferential power of three trajectories per condition.

Graphical Abstract
Three independently seeded 200-ns trajectories were analyzed for FEN-only and FEN+CBD conditions in active-like 5C1M, modeled Morph50 intermediate, and inactive-like 4DKL MOR1 backgrounds. CBD contacts and CBD-associated changes in FEN contacts were receptor-conformation dependent, whereas corrected intracellular TM3–TM6 measures showed no consistent CBD-associated global shift.

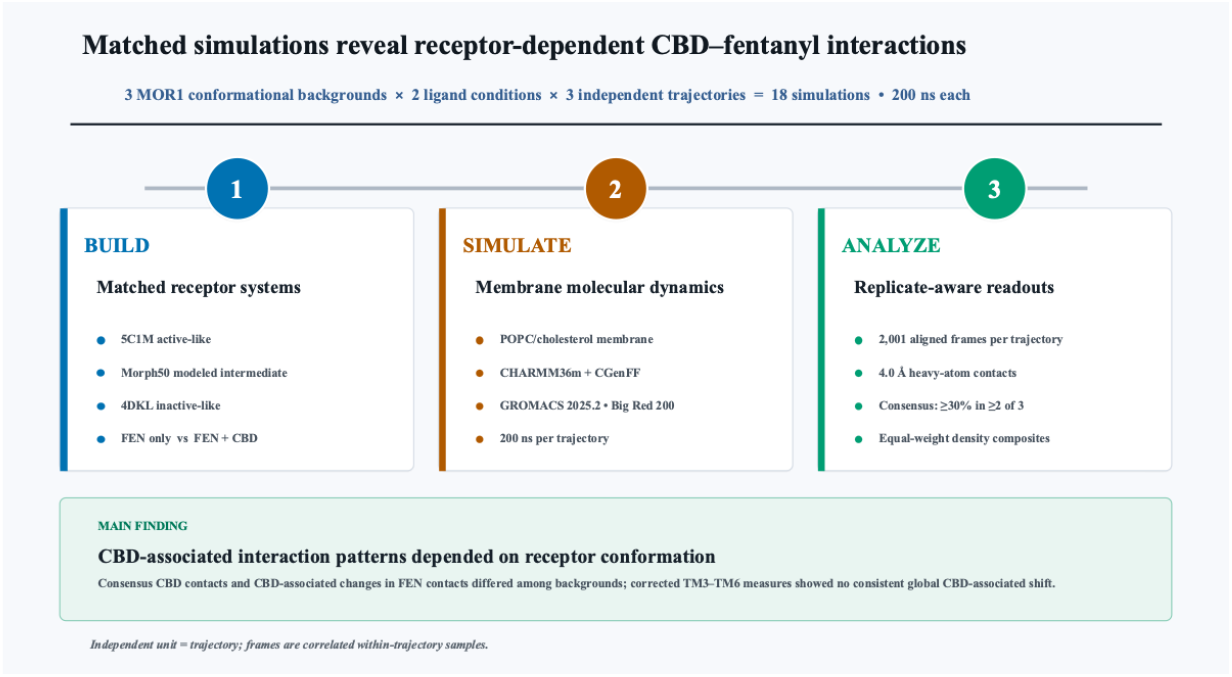

## Scope and study design

This study tests how cannabidiol (CBD), a known μ-opioid receptor 1 (MOR1) allosteric modulator (Bosquez-Berger et al., 2023), is associated with the structural ensemble and fentanyl (FEN) interactions of mouse MOR1 across three receptor backgrounds: an active-like 5C1M-derived model, an inactive-like 4DKL-derived model, and a mouse Morph50 intermediate that we generated along the 5C1M→4DKL transition for the purpose of this study. Each receptor background is simulated with FEN alone and with FEN plus CBD. Three independently seeded trajectories were made per condition, yielding 18 trajectories (*Table I*). All comparisons used the matched 0–200 ns interval from each trajectory. A trajectory, not an individual frame, is the independent replicate. Frames are correlated samples within a trajectory and are used to estimate within-trajectory distributions, densities, contacts, and representative structures.

**Table I.**
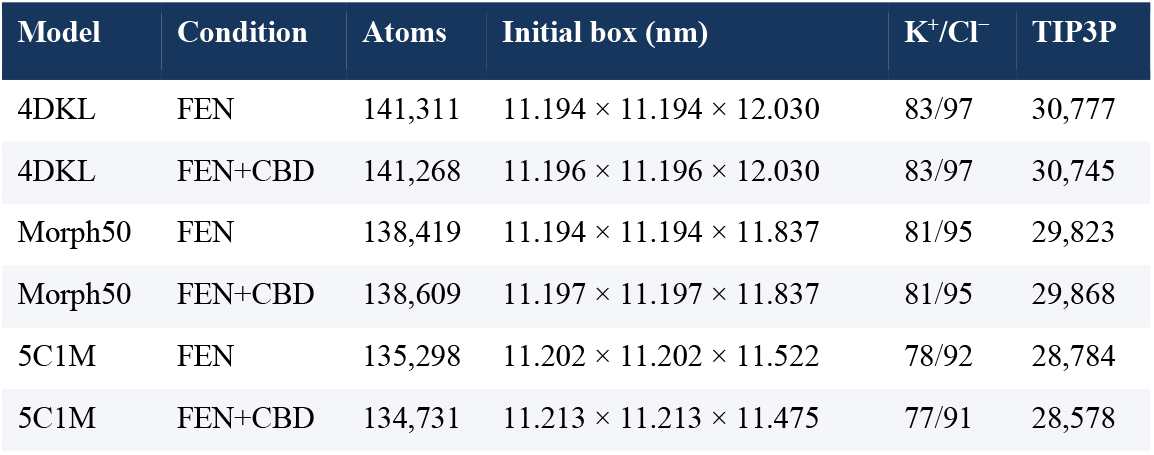
Initial composition and box dimensions of the six membrane-embedded molecular-dynamics systems.

## Methods

### Receptor models

The active-like mouse MOR1 model was derived from PDB 5C1M (Huang et al., 2015). The inactive-like mouse MOR1 model was derived from PDB 4DKL (Manglik et al., 2012). For this model, the T4 lysozyme fusion was removed and the native intracellular-loop 3 segment MLSGSK was reconstructed between the flanking residues R and E, producing the local sequence RMLSGSKE. The repaired model was inspected before ligand placement and membrane construction. The intermediate model was the 50% structure from a corkscrew/linear interpolation (Krebs and Gerstein, 2000) between the mouse 5C1M-derived receptor and the repaired mouse 4DKL-derived receptor. It is therefore termed mouse Morph50 5C1M→4DKL and is treated as a computational intermediate, not an experimentally observed activation state.

### Ligand preparation and placement

Fentanyl and cannabidiol were modeled as noncovalent ligands with residue names FEN and CBD, respectively. CHARMM-GUI Ligand Reader & Modeler (Kim et al., 2017) generated CGenFF/CHARMM-compatible parameters (Vanommeslaeghe et al., 2010). The final coordinate files contain 54 FEN atoms and 53 CBD atoms.

For each receptor background, active 5C1M, intermediate Morph50, and inactive 4DKL, the same initial fentanyl (FEN) pose was used in the FEN-only and FEN+CBD conditions and across all three replicates. Thus, within a given receptor background, the paired conditions differed initially only by the presence or absence of CBD. The initial FEN pose was allowed to differ among 5C1M, Morph50, and 4DKL to accommodate their distinct receptor conformations. Likewise, the initial CBD position was receptor-background specific.

For 5C1M, the FEN pose was transferred by receptor alignment from the fentanyl-bound 8EF5 structure (Zhuang et al., 2022). The CBD pose was obtained by docking to the FEN-containing 8EF5 structure with AutoDock Vina (Eberhardt et al., 2021) and was subsequently transferred with the FEN pose into the 5C1M simulation model; 8EF5 therefore served only as pose provenance and was not included as a replicated simulation comparator. For Morph50 and 4DKL, CBD was docked separately to the corresponding FEN-containing receptor model using receptor-specific search boxes. The selected poses were retained following visual and geometric inspection and incorporated into the respective FEN+CBD starting structures. Consequently, the FEN and CBD coordinates were consistent between conditions and replicates within each receptor background but were not assumed to be identical across the three receptor conformational states.

### Membrane-system construction

Membrane-embedded systems were generated with CHARMM-GUI Membrane Builder (Jo et al., 2008, 2009). Each system contains 114 cholesterol and 266 POPC molecules, TIP3P water, and K+/Cl− ions. Protein, lipid, ligand, water, ion, topology, index, minimization, equilibration, and production inputs are retained in the internal project archive and are not part of the public compact-trajectory release.

### Equilibration and molecular dynamics

Simulations were prepared with the staged minimization/equilibration protocol supplied by CHARMM-GUI and run with GROMACS 2025.2 (Abraham et al., 2015) on Indiana University’s Big Red 200 system. Initial velocities were generated at 303.15 K during step 6.1. Replicate 1 used gen-seed = −1 and its realized seed was recovered from the compiled TPR; replicates 2 and 3 used prespecified unique seeds.

### Matched trajectory processing and alignment

For every run, the 0–200 ns interval is frozen at 100 ps spacing (2,001 frames), centered on the protein, made compact under periodic boundary conditions, and aligned using transmembrane Cα atoms. Model-specific selections preserve homologous transmembrane regions while respecting structure numbering.

**Table II.** GROMACS production molecular-dynamics parameters.

| Setting | Value |
| --- | --- |
| Integrator and timestep | md; 0.002 ps |
| Target production duration | 200 ns |
| Coordinate output | Every 50,000 steps = 100 ps |
| Temperature coupling | v-rescale; SOLU/MEMB/SOLV groups; $\tau_T$ 1 ps; 303.15 K |
| Pressure coupling | C-rescale; semi-isotropic; $\tau_P$ 5 ps; 1 bar; compressibility $4.5 \times 10^{-5} \text{ bar}^{-1}$ |
| Electrostatics | PME; Coulomb cutoff 1.2 nm |
| van der Waals | Force-switch, 1.0–1.2 nm |
| Neighbor list | Verlet; update every 20 steps; 1.2 nm |
| Constraints | Hydrogen bonds constrained with LINCS |

**Table III.** Initial-velocity random seeds used for the three independently seeded trajectories.

| Model | Condition | Replicate 1 TPR seed | Replicate 2 | Replicate 3 |
| --- | --- | --- | --- | --- |
| 4DKL | FEN | 482338719 | 240201 | 340201 |
| 4DKL | FEN+CBD | −1342718467 | 240202 | 340202 |
| Morph50 | FEN | −789005 | 250201 | 350201 |
| Morph50 | FEN+CBD | −73421353 | 250202 | 350202 |
| 5C1M | FEN | 1891628011 | 251201 | 351201 |
| 5C1M | FEN+CBD | 535821007 | 251202 | 351202 |

**Table IV.** Model-specific transmembrane residue ranges used for Cα alignment.

| Model | Transmembrane Cα alignment selection |
| --- | --- |
| 4DKL | resid 4:32, 40:70, 76:106, 118:143, 164:194, 211:242, or 249:262 |
| Morph50 | resid 4:32, 40:70, 76:106, 118:143, 164:194, 211:242, or 249:276 |
| 5C1M | resid 17:45, 53:83, 89:119, 131:156, 177:207, 224:255, or 262:289 |

### RMSD, RMSF, activation, and ligand contacts

Receptor root-mean-square deviation (RMSD) summarizes time-dependent structural displacement after alignment. Per-residue Cα root-mean-square fluctuation (RMSF) summarizes local mobility around each residue’s mean position. Ligand RMSD is calculated from ligand heavy atoms after receptor alignment. The primary activation-related measure was the Cα–Cα distance between the corresponding mouse MOR1 residues Arg165^3.50 on TM3 and Thr279^6.34 on TM6 in each model, using model-specific trajectory residue indices. A sensitivity analysis measured the Cα-centroid distance between intracellular terminal residues 166–170 on TM3 and 275–279 on TM6. Both measures were calculated per trajectory using model-specific structure-number mappings retained in the analysis code.

A protein residue is in contact with a ligand in a frame when any residue heavy atom is within 4.0 Å of any ligand heavy atom. Occupancy is the fraction of the 2,001 frames satisfying this criterion within one trajectory. Mouse MOR numbering is reported alongside structure numbering using the validated +64 offset for 4DKL/Morph50 and +51 offset for 5C1M.

### Ensemble density and representative frames

Ligand density maps were calculated after model-specific transmembrane alignment on a shared 1 Å grid. Each replicate density was normalized separately and smoothed with a Gaussian kernel (σ = 1 voxel) before the three maps were combined by arithmetic mean, giving each independent trajectory equal weight. Publication surfaces independently enclose the highest-density 50% of each ligand distribution. A representative ligand position was selected from the trajectories based on how well it fit within the composite density map. It is shown only to help orient the density envelope and should not be interpreted as a unique or preferred binding pose.

The receptor wireframes used for structural context are the aligned model-specific replicate-1 FEN-only receptors at 0 ns. Receptor views were oriented extracellular side up, with the amino-terminal end at the top and the carboxyl-terminal end at the bottom.

### Secondary pocket-geometry analysis

As a secondary structural-context analysis, fpocket 4.0 (Le Guilloux et al., 2009) was run independently on receptor-only dominant-cluster representative structures from the FEN-only and FEN+CBD conditions. Ligand-associated pocket components were selected using fpocket alpha-sphere vertices within 4.0 Å of ligand heavy atoms. When a pocket component approached both ligands, its vertices were assigned to the nearer ligand rather than duplicated. This static geometric analysis was used for supplementary visualization and quality control only; it did not define the trajectory-wide ligand contacts or composite density maps and was not treated as an independent replicate.

### Replicate-level analysis

Before examining replicate 3, we prespecified a consensus interacting residue as one with a contact occupancy of ≥30% in at least two of the three replicate trajectories for a given receptor background and ligand condition.

The trajectory was the unit of replication; individual simulation frames were not treated as independent observations. Results are therefore presented for each replicate trajectory together with the equal-weight mean across replicates and the observed between-trajectory variability. No frame-level *P* values were calculated.

For each receptor background, the effect of CBD was calculated within matched replicates as the FEN+CBD value minus the corresponding FEN-only value. These three replicate-level contrasts were then summarized by their magnitude and directional agreement. With only three replicate contrasts, formal inference is necessarily limited. We therefore report the contrasts descriptively and do not use them to make population-level claims of statistical significance.

### Software and reproducibility

Production simulations were performed using GROMACS 2025.2 (Abraham et al., 2025; Abraham et al., 2015). The final replicate analysis used MDAnalysis 2.10.0 (Gowers et al., 2016; Michaud-Agrawal et al., 2011), NumPy 2.3.2 (Harris et al., 2020), and SciPy 1.16.1 (Virtanen et al., 2020); molecular graphics were prepared with UCSF ChimeraX 1.11 (Pettersen et al., 2021). System preparation used AutoDock Vina (Eberhardt et al., 2021; Trott & Olson, 2010) for ligand docking, CHARMM-GUI Membrane Builder (Wu et al., 2014) and Ligand Reader & Modeler (Kim et al., 2017), CGenFF (Vanommeslaeghe et al., 2012) compatible ligand parameters, and Modeller (Šali and Blundell, 1993) for the 4DKL loop reconstruction. Secondary pocket-geometry analysis used fpocket 4.0 (Le Guilloux et al., 2009). The public deposit includes compact trajectories with matching starting-coordinate structures, machine-readable analysis outputs, locked software versions, analysis scripts, provenance, and validation records; the companion publication-results archive contains the figure-generation scripts and final figures. During preparation of this manuscript, the authors used OpenAI Codex with the GPT-5.6 Sol model at medium reasoning effort to assist with computational workflow execution, file organization, and language editing. All commands, analytical outputs, and manuscript changes were reviewed by the authors, who take full responsibility for the final work.

## Results

### Trajectory integrity and replicate-level metrics

All 18 compact trajectories passed quality-control checks for frame coverage and atom counts. Each trajectory contained 2,001 frames sampled at 0.1-ns intervals from 0.0 through 200.0 ns. All numerical arrays contained finite values, and all density values were nonnegative. Raw ligand-density grids were normalized to unit integrated probability by dividing the voxel counts by the total number of ligand heavy-atom observations and the 1 Å^3^ voxel volume. Gaussian smoothing preserved the normalized total probability, apart from negligible numerical rounding. Contact tables contained no duplicate records for the same receptor background, condition, replicate, ligand, protein segment, residue number, and residue name. Reanalysis of unchanged trajectories exactly reproduced the corresponding archived per-frame metrics, replicate summaries, residue-contact occupancies, and Cα RMSF values. Within each receptor background, all ligand-density maps were constructed on a common grid spanning all conditions and replicates.

The equal-weight mean Arg165^3.50^–Thr279^6.34^ Cα distance was 6.59 Å with FEN alone and 6.51 Å with FEN+CBD in 4DKL, 9.52 and 9.72 Å, respectively, in mouse Morph50, and 12.71 and 12.56 Å, respectively, in 5C1M. The within-replicate mean changes (FEN+CBD minus FEN) were −0.08, +0.20, and −0.14 Å, and no receptor background showed the same direction in all three independent replicates. As a secondary geometric measure, we calculated the distance between the Cα centroids of intracellular TM3 residues 166–170 and TM6 residues 275–279. The equal-weight mean decreased from 11.57 to 10.72 Å in 4DKL and from 15.18 to 14.68 Å in 5C1M in all three paired replicates. In Morph50, the centroid mean decreased from 12.88 to 11.78 Å, but the direction was inconsistent and the mean was driven by a large decrease in replicate 2. Thus, the apparent CBD-associated change depended on the geometric definition.

FEN RMSD relative to each trajectory’s aligned initial pose showed the greatest between-replicate variability in 4DKL. Mean FEN RMSD values for the FEN-only trajectories were 2.21, 6.99, and 3.18 Å for replicates 1–3, respectively, compared with 4.15, 4.47, and 3.48 Å for the corresponding FEN+CBD trajectories. The elevated FEN RMSD in FEN-only replicate 2 was not observed in replicates 1 or 3, reflecting replicate-specific trajectory heterogeneity. Between-replicate variation in FEN RMSD was smaller for 5C1M and Morph50.

### State-dependent consensus CBD contacts

Using the prespecified consensus rule of at least 30% contact occupancy in at least two of three independent FEN+CBD trajectories, 9 CBD-contact residues were identified in 4DKL, 11 in mouse Morph50, and 8 in 5C1M (Figure 1). The contact patterns differed among receptor backgrounds. In 4DKL, the largest equal-weight mean occupancies occurred at TYR128, ASN127, GLN124, TRP133, and GLY131. Mouse Morph50 showed high occupancy at TYR148, LEU219, LYS233, and TRP318, whereas 5C1M was dominated by CYS217 with additional contacts including TRP318, LEU219, and ILE144.

**Figure 1.**
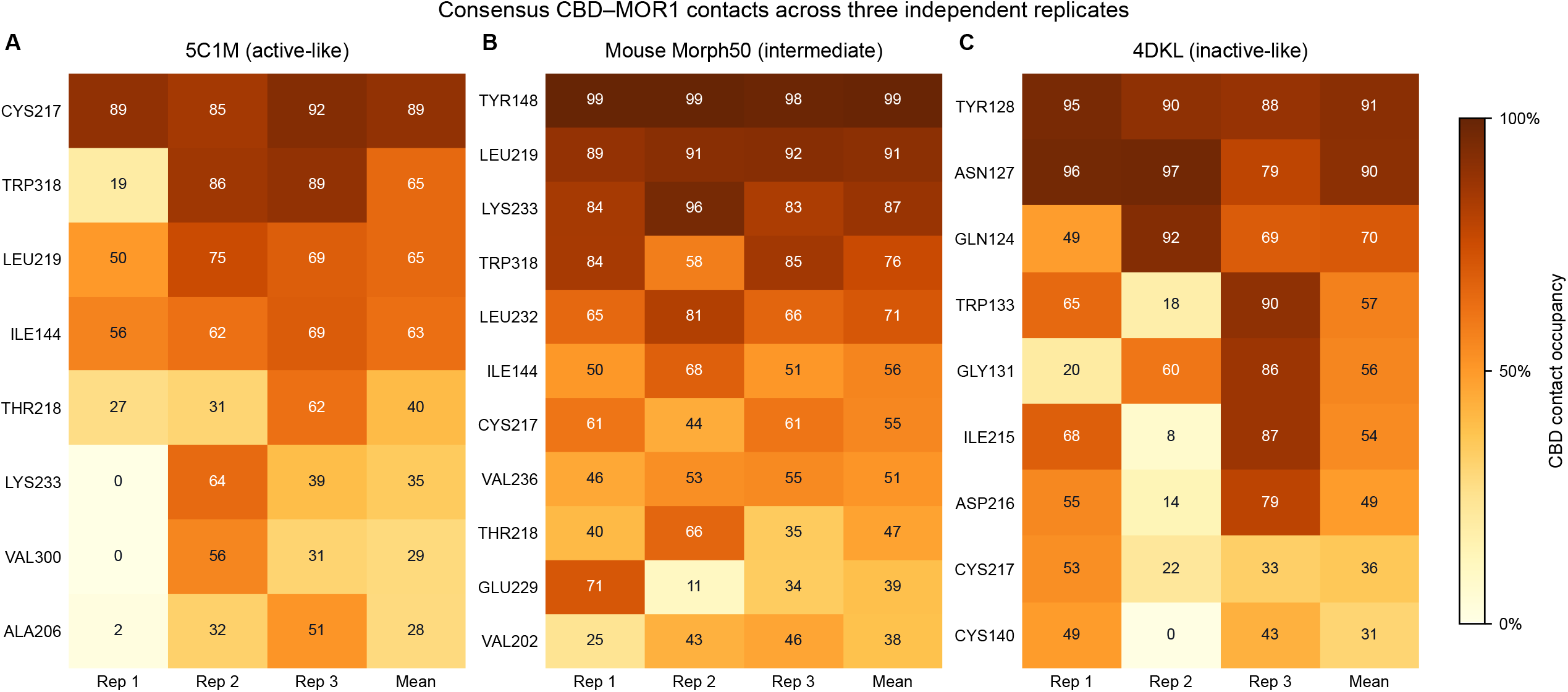
Consensus CBD-MOR1 contacts across three independent replicates. CBD-residue heavy-atom contact occupancy over the matched 0-200 ns window for (A) 5C1M active MOR1, (B) mouse Morph50 5C1M-to-4DKL intermediate MOR1, and (C) 4DKL inactive MOR1. A residue is included when CBD contact occupancy is at least 30% in at least two of three independent FEN+CBD trajectories. Heat-map values are percentages for each replicate, and the equal-weight replicate mean. A mean can be below 30% when two replicates meet the threshold and the third has low occupancy. Contact was defined by any ligand-residue heavy-atom distance of 4.0 A or less in a frame.

### CBD-associated changes in FEN contacts

Within-replicate FEN+CBD-minus-FEN comparisons identified several contact changes with the same sign in all three trajectories (Figure 2). In 4DKL, the largest mean increases were observed at TRP318 (+39 percentage points), VAL300 (+38), and ILE322 (+25). Mouse Morph50 showed a pronounced decrease at TRP293 (−53) and increases at MET151 (+28) and ILE296 (+17). In 5C1M, TRP318 increased (+35), whereas HIS297 (−17) and CYS140 (−14) decreased. Directional concordance across three trajectories is descriptive and does not constitute a formal significance test.

**Figure 2.**
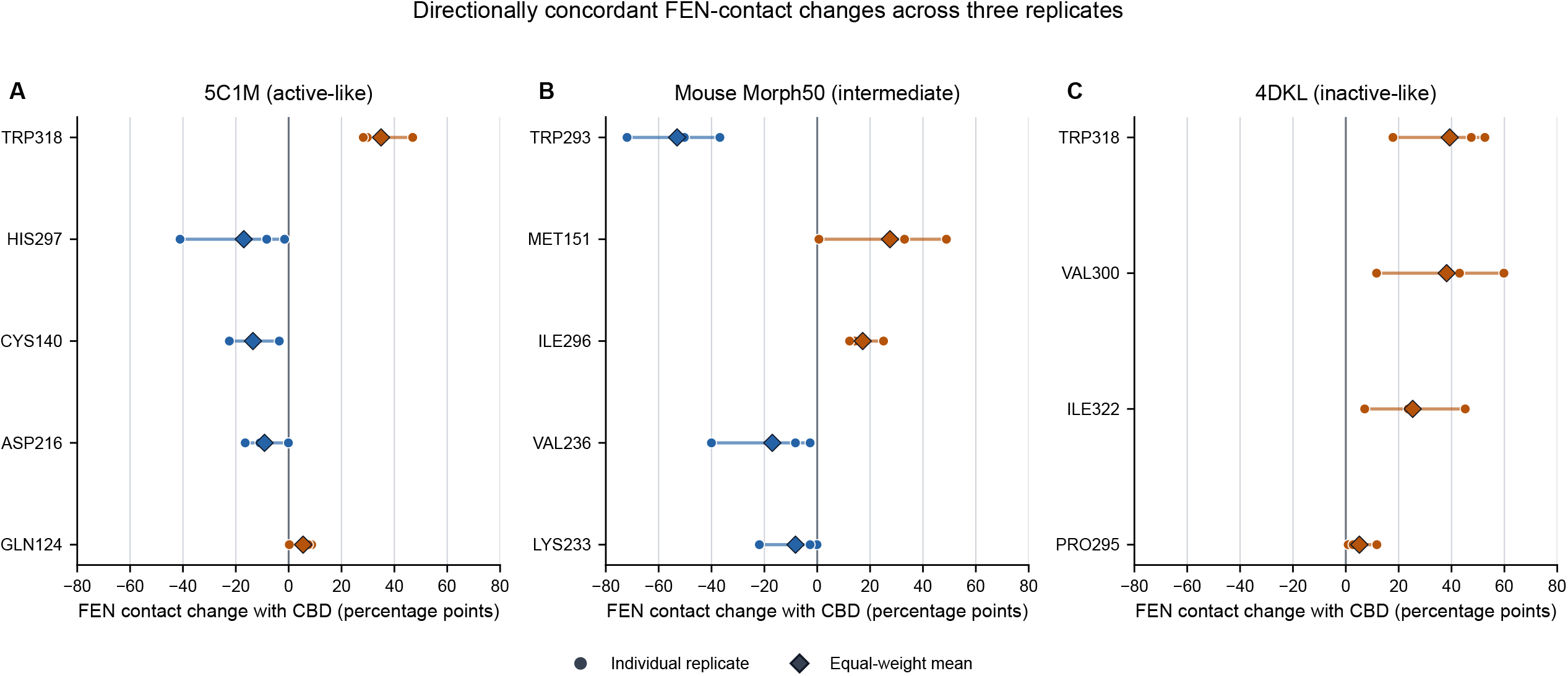
Directionally concordant FEN-contact changes with CBD. Within-replicate changes in FEN-residue contact occupancy for (A) 5C1M, (B) mouse Morph50, and (C) 4DKL. For each replicate, delta occupancy is FEN occupancy in the FEN+CBD trajectory minus FEN occupancy in the corresponding FEN-only trajectory. Circles represent individual replicate values, horizontal segments show the replicate range, and diamonds show equal-weight replicate means. Positive values indicate greater FEN contact occupancy when CBD is present; negative values indicate lower occupancy. Displayed residues have the same nonzero direction in all three replicates and an absolute mean change of at least five percentage points. Directional concordance is descriptive and is not a formal significance test.

### Replicate-composite ligand distributions

Equal-weight composite density maps demonstrated distinct ligand distributions across the inactive-like, modeled intermediate, and active-like receptor backgrounds (Figure 3). Separate normalization prevented a trajectory from dominating because of frame count, while averaging retained regions sampled reproducibly across replicates. The broader 4DKL FEN distribution was consistent with its larger between-trajectory RMSD heterogeneity. Representative FEN and CBD poses are displayed only to orient the density envelopes and are not interpreted as uniquely preferred structures.

**Figure 3.**
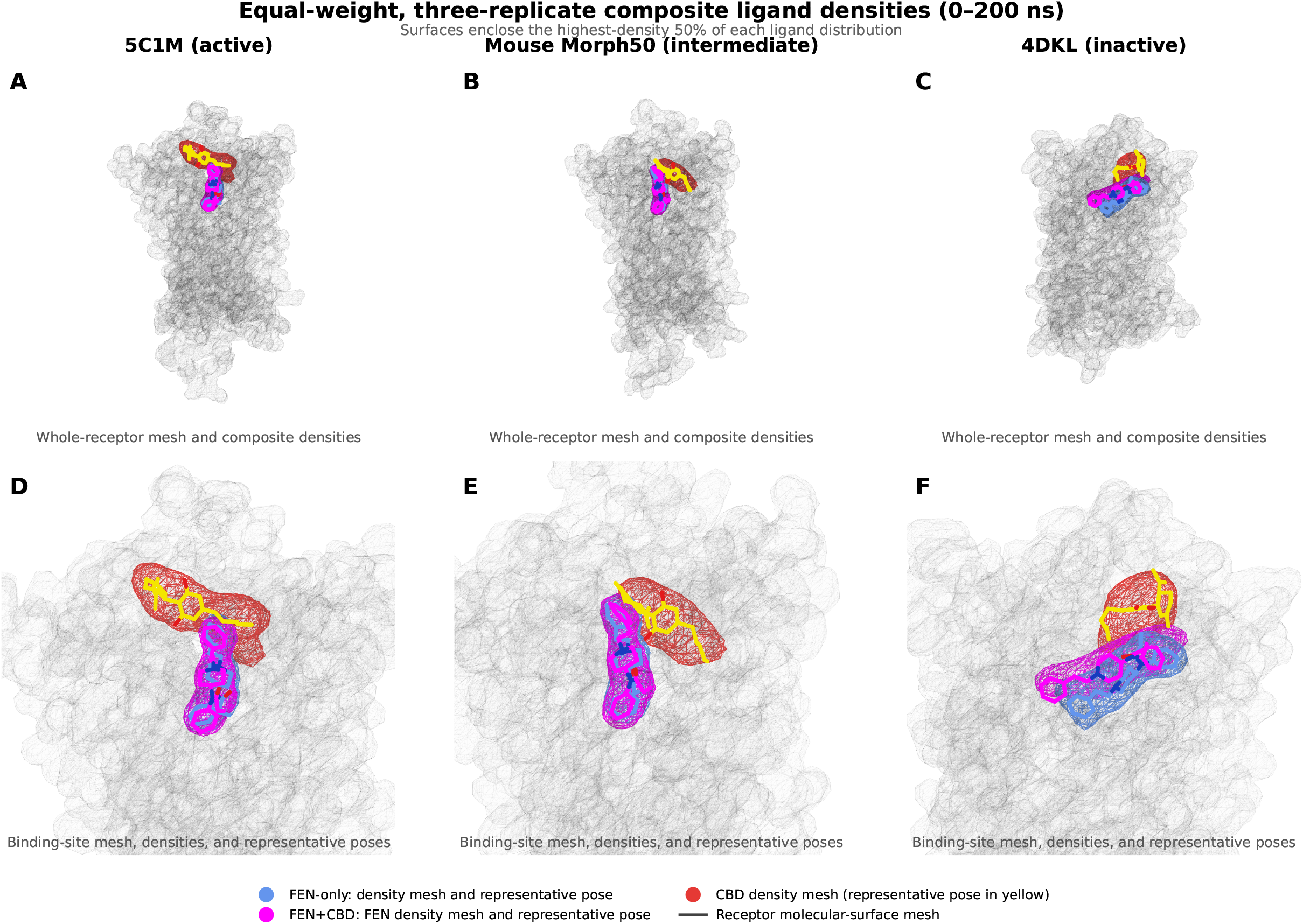
Three-replicate composite ligand-density distributions. Equal-weight replicate-composite ligand densities over the matched 0-200 ns window for 5C1M active MOR1 (A, D), mouse Morph50 intermediate (B, E), and 4DKL inactive MOR1 (C, F), shown in whole-receptor context (A-C) and as binding-site close-ups (D-F). Cornflower blue denotes the FEN distribution in FEN-only trajectories, magenta denotes the FEN distribution in FEN+CBD trajectories, and red denotes the CBD distribution in FEN+CBD trajectories. Each replicate map was normalized by its own ligand heavy-atom observations before the three maps were combined by arithmetic mean. Maps use a 1 A grid and Gaussian smoothing with sigma = 1 voxel. Each surface independently encloses the highest-density 50% of its distribution. The black/gray wireframe is the aligned model-specific replicate-1 FEN-only receptor at 0 ns. Cornflower-blue and magenta sticks show representative FEN poses, and yellow sticks show the representative CBD pose. For each density map, we selected the sampled ligand pose that best matched the regions most consistently occupied across the three equally weighted replicates. The displayed pose is provided only as a visual reference for interpreting the density envelope. Receptor views use an extracellular-up orientation, with the amino-terminal end at the top and the carboxyl-terminal end at the bottom.

## Discussion

### Central interpretation

The principal finding is that CBD-MOR1 contacts and CBD-associated changes in FEN contacts were receptor-state dependent rather than uniform across the three modeled backgrounds. The active-like 5C1M, modeled Morph50 intermediate, and inactive-like 4DKL systems displayed different CBD-contact neighborhoods and different directionally concordant FEN-contact changes. In contrast, the corrected intracellular TM3–TM6 measures did not provide consistent evidence that CBD shifted each receptor background toward a common global activation state. These observations support an ensemble-based interpretation in which CBD co-presence is associated with locally distinct FEN-MOR1 interaction patterns in different receptor conformations.

The functional cAMP findings of Bosquez-Berger et al. (2023) support negative-allosteric-modulator activity of CBD and several CBD analogs, whereas preferential stabilization of an inactive MOR1 conformation was proposed from comparative docking as a possible mechanism. Our simulations do not contradict the functional inhibition but qualify this structural interpretation. CBD did not produce a replicate-consistent change in the primary Arg165^3.50–Thr279^6.34 activation-related coordinate, and the terminal-centroid contractions observed in 5C1M and 4DKL were not reproduced uniformly across geometric definitions or in Morph50.

Even sustained CBD association with MOR1 would not necessarily cause a large TM3–TM6 displacement, because ligand binding, allosteric efficacy, and sampling of receptor conformations are distinct properties. A transition from the active-like 5C1M background toward an inactive-like 4DKL conformation could, in principle, produce a contraction of several ångströms along the Arg165^3.50–Thr279^6.34 coordinate. However, each trajectory sampled only 200 ns continuously; combining independent trajectories increases replication but does not create a single millisecond-scale simulation or ensure that slow conformational barriers will be crossed. The absence of a reproducible several-ångström shift therefore indicates that a large CBD-associated transition was not sampled under these conditions, rather than demonstrating weak CBD binding or an absence of negative allosteric activity. CBD-associated inhibition could instead involve localized changes in helix geometry, microswitch coupling, ligand efficacy, or receptor conformational populations without producing a complete transition to the crystallographic inactive state.

### Replicate heterogeneity and limitations

The completed 4DKL replicate 3 analysis was important because it did not reproduce the unusually large FEN displacement seen in FEN-only replicate 2. Reporting all replicate values and equal-weight density composites prevents that trajectory from being hidden by a pooled-frame analysis or, conversely, from being treated as the sole behavior of the inactive-like model. Between-trajectory heterogeneity remains a biological and computational feature of the sampled ensemble rather than an error to be averaged away.

This study is limited by three independent trajectories per condition, the finite 200 ns window, starting-pose dependence, force-field approximations, and the computational nature of the Morph50 intermediate. The simulations describe mouse receptor models, while human 8EF5 was used only as provenance for transfer of the 5C1M FEN pose and was not a replicated comparator. Frames within trajectories are correlated, and no frame-level inferential tests were performed. The findings therefore generate specific state- and residue-level hypotheses that require experimental validation.

## Data availability

The compact receptor-plus-ligand trajectory subsets, matching 0 ns reference structures, contact and trajectory-metric tables, Cα RMSF results, replicate-specific and equal-weight ligand-density maps, validation records, analysis scripts, and publication source data are deposited in Zenodo [10.5281/zenodo.22050916]. The release contains 18 matched 0-200 ns compact trajectories but does not include the full lipid, solvent, ion, topology, checkpoint, or production-input collection required to restart the original simulations. Data, figures, tables, and documentation are released under CC BY 4.0; original analysis and figure-generation scripts are released under the MIT License.

